# A practical bacterial biodosimetry procedure to assess performance of lab-scale flow-through ultraviolet water disinfection reactors

**DOI:** 10.1101/2022.12.23.521800

**Authors:** Philipp Sperle, Mohammad S. Khan, Jörg E. Drewes, Christian Wurzbacher

## Abstract

Biodosimetry can be used to estimate the fluence of a reactor by determining its ability to inactivate a challenge organism. Especially for small-scale flow-through reactors, inconsistent procedures are reported for bacterial cells. This study aims to develop a standardized, simple procedure for bacterial biodosimetry in flow-through UV systems with relevant biofilm forming bacteria in order to evaluate biofouling control by UV. In particular, the challenge organism, the type of experimental setup and the execution of single steps during biodosimetry with bacterial cells can cause largely deviating results. Since previous work was restricted to model organism, which are not relevant for biofouling, we critically reevaluated all reported steps for the biofilm forming *Aquabacterium citratiphilum* and identified three main factors for biodosimetry reproducibility in flow-through systems: Protractions of cells from negative controls can heavily impact inactivation efficacy, but can be reduced by ordering samples by decreasing fluence. Further, to avoid photo repair, samples must be processed under red light only. Lastly, biofilm forming bacteria can strongly absorb on plastic labware, which requires counter measures in form of special labware and the addition of surfactants. Overall, the developed protocol provides a biodosimetry standardization for bacterial cells of flow-through systems, facilitating reproducibility and transferability of results between studies that use bacterial cells as challenge organism.

**Synopsis:** Biodosimetry is used to characterize performance of flow-through ultraviolet reactors, but artifacts can occur if not addressed properly.

## 1. Introduction

Using ultraviolet (UV) light as a disinfection process has been researched for more than 50 years and is a well-established technology (*1*). UV disinfection has the great advantage of causing limited by-product formation and being chemical-free (*2–4*). In general, UV light can be categorized according to its applied wavelength with UV-A covering 315-400 nm, UV-B 280-315 nm, and UV-C 200-280 nm. Even before unraveling the exact impact of UV light on DNA, it was elucidated that DNA is mainly involved as absorbing material for UV light leading to the inactivation of microorganisms through the usage of action spectra (comparing the relative efficiency of incident light vs wavelength) (*1*). Several photoproducts can be formed due to UV irradiation, however, cyclobutene-pyrimidine dimers and 6-4 photoproducts are considered to be of main importance for DNA damage (*1, 5*). Besides, for longer wavelengths like UV-B, the production of reactive oxygen species can play a considerable role in indirect damage to the cell’s components, proteins, and DNA (*6*). Nevertheless, UV-induced damage might not necessarily be lethal and bacterial cells might be capable to recover due to dark or photo repair mechanisms by light in the range of 310 to 480 nm (*1*). Especially for bacteria, overall disinfection efficiency can be heavily influenced by photo repair (*7*).

Even though some studies observed deviations (*8, 9*), UV disinfection is commonly assumed to follow the Bunsen–Roscoe reciprocity law, meaning UV disinfection solely depends on the product of fluence rate and time (fluence). Following the one-hit-one-target function, logarithmic survival behaves linearly over fluence (*1*). However, according to Harm (*1*), this inactivation behavior might not be observed over the complete fluence range. At low fluencies, curves with shoulders might be observed as the inactivation rather follows a multi-target or multi-hit model or due to repair processes being more efficient at lower fluence. In addition, upward concave curves can be observed if multi-components (multiple fractions with different inactivation rates) are present. In higher fluence regions, inactivation curves might show tailing behavior, likely caused by UV resistant subpopulations, reactor hydraulics, cell aggregates or other shielding effects (*10–14*).

Common sources for UV irradiation include low and medium pressure (LP and MP), and xenon flash lamps. In recent years UV-LEDs received increasing attention as they have advantages such as not containing mercury, being robust with fast start-up times, and potentially being more energy efficient with a longer lifetime (*13*). Furthermore, according to Song et al. (*15*) they are compact as well as available in different wavelengths and viewing angles which enables new reactor designs and novel applications. Hence, when designing new reactors especially hydrodynamics, radiation distribution, and kinetics require consideration and evaluation. In general, lab-scale systems not only deviate from the scale of application but also new reactor designs or treatment combinations are tested using different organisms for challenge tests. Especially through their compactness (*15*), the newly available LEDs allow building of small-scale reactors with variable designs.

There are several methods available to quantify fluence in UV reactors (*16*), but the common method for certifying a new reactor is biodosimetry (*17, 18*). Biodosimetry cannot estimate fluence distribution directly but it directly measures the inactivation of microorganisms (*16*), being the primary goal in water disinfection. The concept was first applied by Qualls and Johnson (*19*). In this paper and according to the U.S. EPA (*17*), biodosimetry is referred to as the experimental test during the validation procedure, in which the logarithmic inactivation of a challenge organism is tested under desired flow rates, UV transmissions (UVT), and lamp power combinations. In the next step, the results from the biodosimetry test are then compared to a dose-response curve of the same challenge organism solution recorded under controlled conditions usually using a collimated beam apparatus (CBA) to calculate the reduction equivalent dose (RED) and inactivation credits. In general, the UV disinfection capability of a flow-through reactor might be lower compared to a CBA. Ideally all microorganisms should receive the same fluence, but in practice due to hydraulic shortcuts or similar imperfections, a fraction of microorganism might exist receiving less fluence influencing the disinfection efficacy (*17*). For reactor validation, commonly used challenge organisms include MS2 phage and endospores of *Bacillus subtilis*. For both targets, standardized methods for large-scale and batch reactors (CBA) are available. In contrast, a general standardize procedure for vegetative microbial cells is currently lacking. Microbial cells can have diverse cell wall components with different capabilities for photon adsorption through heterogeneous pigments, and in contrast to phages and spores, cells may actively respond to radiation by photo repair mechanism (*1, 5*). If the microorganism used for validation shows a different UV sensitivity than the target organism, RED needs correction for so called RED bias, depending on UV sensitivity and fluence distribution of the UV reactor (*14*). Furthermore, differently applied wavelengths lead to different disinfection efficiencies that must be considered for each microorganism (*20*).

As a consequence, a range of different methods for biodosimetry using microbial cells as challenge organism have been applied to lab-scale reactors (*3, 21–29*). However, the parameters for these vary and are not well defined, ranging from flow-through to batch systems with or without recirculation, a range of species in different conditions and feed solutions, varying washing steps and with or without inclusion of photo repair of microorganism (*3, 21–29*). There are many steps involved in biodosimetry that can be critical, including cultivation, washing, the design of the experiment, and processing of the samples including measures for or against repair mechanisms(*3, 21–29*). However, as the methodology is not yet standardized, results may largely differ between studies. Furthermore, most studies focused on nosocomial bacteria or model organism such as *Escherichia coli* (*3, 21–29*), which are not relevant in technical applications that are affected by biofouling. Biofouling of membranes is caused by biofilm-forming, usually non-infectious, non-model organism, which to our knowledge have not been used as challenge organism, yet.

This study aims to provide a standardized biodosimetry protocol for flow-through reactors and to elucidate critical steps that mitigate artifacts exploring new UV treatment designs with bacterial cells (using LEDs or conventional lamps). This is achieved by comparing the existing methods for flow-through UV reactors and the subsequent evaluation of the most important steps resulting in a standardized and reproducible biodosimetry protocol for the non-model organism, *Aquabacterium citratiphilum*, a gram-negative bacterium found typically in drinking water biofilms around Europe (*30*) with potential relevance for biofouling of high-pressure membrane treatment (*31*).

## 2. Material and Methods

### 2.1. Testing skid for biodosimetry experiments

The testing skid to perform biodosimetry experiemnts (Figure 1) consisted of a feed container that was continuously stirred with a magnetic stirrer (MR 2000, Heidolph, Germany). Feed was withdrawn using the gear pump (DGS.38PPPV2NN00000, Tuthill Alsip, United States and K21R 63 K2 H, VEM motors Thum, Germany) and pump speed was regulated using the frequency converter (FUS 037/E2, PETER electronic GmbH & Co. KG, Germany). The flow was measured using a magnetic-inductive flow meter (MIK-6FC08AC34P, KOBOLD Messring GmbH, Germany). The flow of the pump was controlled over a PI controller implemented in the computer-aided control system. A UV-LED reactor was placed in a temperature cabinet (Innova 4230, New Brunswick Scientific, United States) at 15°C and the reactor was directed in an upright position to avoid any air bubble accumulation. After passing through the LED reactor, the effluent was directed to the waste container, which can be switched with sampling vials. Pipes between the different elements had an inner diameter of 4 mm and were made of black PTFE. Only the last approximately 50 cm were made from a transparent PTFE pipe with an inner diameter of approximately 1 mm to provide more resistance for the water and therefore higher process stability at very low flows. The pipe was covered with aluminum foil to reduce light from the surrounding.

**Figure 1.**
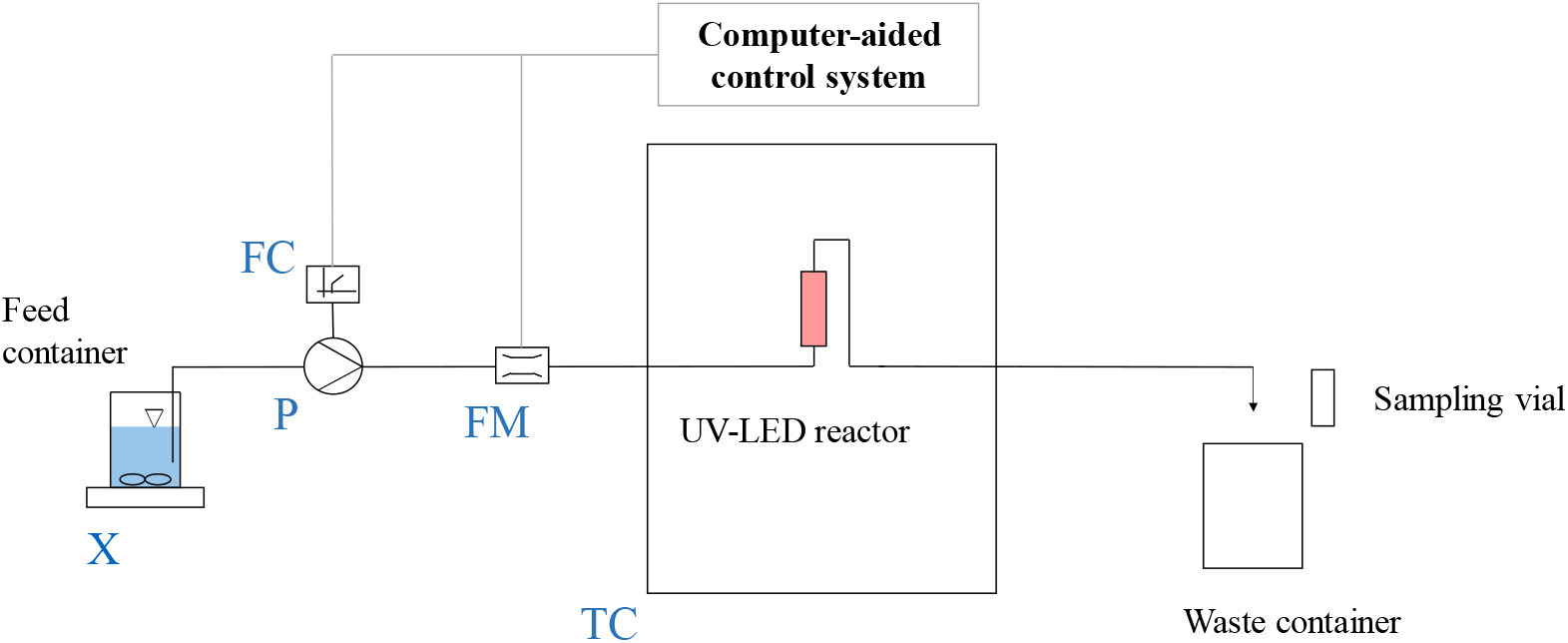
Schematic of the employed laboratory skid for biodosimetry experiments; X = magnetic stirrer, P = gear pump, FC = frequency converter, FM = flow meter, TC = temperature cabinet

Our simplified UV-LED reactor was provided by UV-EL GmbH (Dresden, Germany) and consisted of a reflective PTFE block as housing, which covered the LEDs and the silica glass pipe through which the water was flowing. Silica glass pipe had a total length of 10 cm and an inner diameter of 0.7 cm. The central6 cm of the 10 cm glass pipe were covered with PTFE and are seen as the reactor volume. Over the LEDs, the LED cooling system, consisting of a metallic cooling body and a fan, was attached to the PTFE block and the LEDs themselves. As LED module, the 2P2S-S6060 UVC Quad SMD Module (Bolb Inc., United States) consisting of four LEDs with a peak wavelength between 265 to 275 nm was used (*32*). A schematic drawing is presented in the supplementary materials (Figure S1). The LED was powered using the LED-Pulse-Controller by Leistungselektronik JENA GmbH (Germany). The pulse controller enabled flexible change of the LED output power by adapting the driving current in steps of 8 mA. In general, the higher the driving current, the higher the fluence rate. Whereas fluence rate of the system depends on the driving current of the LED, fluence depends on fluence rate multiplied by hydraulic retention time (HRT). The HRT within the experiments was varied by adapting the flowrate within the system and the HRT was calculated by dividing reactor volume through flowrate. HRT was varied in equidistant steps. After switching to a different HRT or irradiation setting, a fixed volume was flushed through the system based on a tracer test to reach a steady state in the outflow.

### 2.2. Tracer test

To characterize the hydraulics and the mean hydraulic retention time within the skid (Figure 1), a simple positive step tracer test tracking electric conductivity (EC) at two flow rates (1.5 and 6.0 L h^-1^) was performed. For this purpose, the skid was flushed with DI water and then the feed was switched to local tap water. A flow cell with a conductivity meter attached to the outlet monitored the outflow EC until it became steady and indifferentiable from the feed averaging across 1 sec time intervals. As EC at the beginning of the experiments was close to 0, the initial EC value was subtracted from the EC readings. Analogous to Levenspiel (*33*) the cumulative residence time distribution (RTD) curve F(t) was calculated:

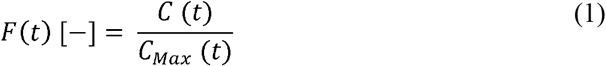

Whereas conductivity values were used as a surrogate for total dissolved solids concentration. By differentiating F(t) for time t, the residence time distribution E(t) was obtained (*33*).

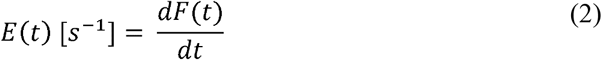

With knowledge about E(t), mean residence time t_m_ and variance σ^2^ were calculated (*34*):

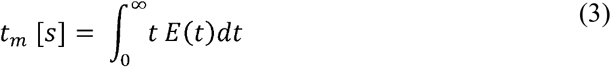

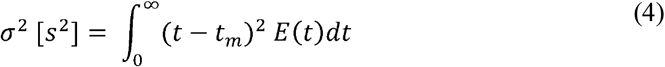

Space-time τ was measured by draining the whole system by first running the pump without feed and then flushing the system with pressurized air. The drained water was weighed and τ calculated by dividing the volume of water through flow (*33*). To compare tracer results to other reactors normalized resident times were calculated based on Fogler (*34*).

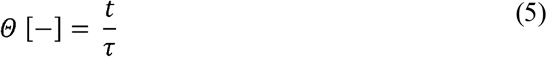

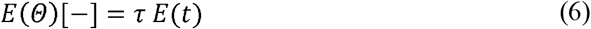

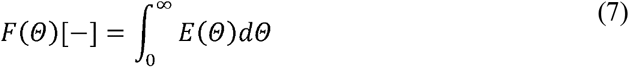

Recorded cumulative RTDs were further compared to two idealized models, a tank in series and a laminar flow model.

The tanks in series model was fitted with the number of tanks n in series as described by (*34*).

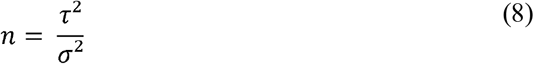

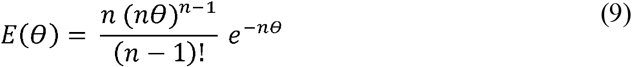

The laminar flow model was calculated according to Fogler (*34*).

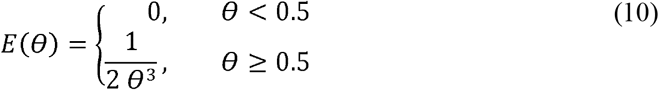

### 2.3. Preparation of *Aquabacterium citratiphilum*

*Aquabacterium citratiphilum* (DSM 11900) was obtained from the Leibniz Institute DSMZ-German Collection of Microorganisms and Cell Cultures GmbH (Germany), grown on suitable agar plates or bouillon with Medium 830a (*35, 36*). For the cultivation of *A. citratiphilum* in liquid and establishing a growth curve, a colony grown on an agar plate was transferred into the liquid bouillon in a 250 mL Erlenmeyer flask, which was shaken at 120 rpm at 20°C (*30*). To calibrate a grow curve with optical density (OD, 600 nm), the cell suspension was plated and colony forming units (CFU) were counted using a cell density meter WPA CO 8000 (Biochrom, United States) and colony counter BZG 30 (WTW, Germany) (Figure S2).

### 1.1. Experimental procedure

For biodosimetry feed solutions, cells with an OD between 0.24 and 0.54 in exponential growth phase were harvested by a centrifugation step at 5,000 RCF for 10 min at 4°C. The supernatant was withdrawn and the pellet dissolved in phosphate-buffered saline (PBS), prepared according to NSF/ANSI 55 (*18*). This washing step was repeated three times in total. Based on the growth curve and measured OD, appropriate amount of washed cell suspension was used to reach a final concentration of approximately 5 ·10^6^ CFU mL^-1^ in 2.5 L PBS. For reducing cell adsorption, PBS was modified with Tween20 (0.002%), when needed. Tween20 was purchased from Sigma-Aldrich, France). After washing, the feed solution should be used immediately to avoid cells from adapting to the new low nutrient conditions (*37*). For the final three experiments as shown in Chapter 3.4, by using OD of bouillon, cell density of 5.9 · 10^6^ mL^-1^ with a standard deviation of 10% was achieved. Through washing, the decadic absorption coefficient at 273 nm was maintained at 0.043 ± 0.014 cm^-1^. In general absorbance should be adapted to desired value through the addition of additives (*17*). In case lower absorbance would be needed, cell number in the feed could be reduced. In this study absorbance was not further changed.

Before starting an experiment, the experimental skid was flushed with DI water, followed by a disinfection through recirculating 250 mL of 1% H_2_O_2_ (Merck, Germany) for 10 minutes, but discarding the first 100 mL to avoid H_2_O_2_ dilution. Then, the system was primed with 1 L sterile MilliQ, followed by 1 L of sterile PBS. During priming, the LED was activated for 10 minutes. The feed container was connected to the inlet and a waste container was placed below the outlet, where the samples were taken. Samples for different UV hydraulic retention times were taken in triplicates and random order in 15 mL transparent tubes. Blank samples (no UV irradiation) were taken at the same spot, before the first irradiation sample, after the second, and after the last sample. Afterward, samples were diluted serially using PBS to reach 25-250 CFU per plate analogous to (*18*). 0.1 or 0.15 mL, depending on the expected CFU, was pipetted on room temperature agar plates and spread using a Drigalski spatula made of glass (height 200 mm, width 45 mm and diameter 4 mm) till a resistance between plate and spatula could be felt. Spatula was dipped into 80 % ethanol and flamed. Overall, three identical spatulas were used alternately so each spatula had sufficient time to cool down. Plates were inverted and inoculated in dark at 20°C for 7 days. CFU were counted using the BZG 30 (WTW, Germany). The log removal value (LRV) was calculated following equation 11:

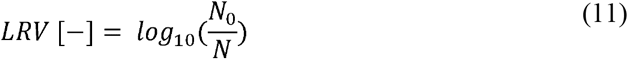

Whereas N_0_ represents the geometrical mean of the unirradiated samples in CFU mL^-1^ and N is the geometrical mean of an irradiated sample at a certain HRT (*18*). Bouillon as well as PBS used for washing and dilution were plated as negative controls.

## 3. Results and Discussion

Within the past years, a variety of biodosimetry studies were conducted at pilot- and lab-scale for flow-through reactors using bacteria as challenge organisms. Our literature search (August, 2022) on bacterial biodosimetry experiments of flow-through reactors resulted in ten studies (excluding experiments with spores; including two studies for batch operation) that shared some steps of a biodosimetry experiment (as washing steps, cultivation and processing of samples). Overall, the different studies investigated revealed major differences and no standardization across many of the experimental parameters (like growth phase, cell washing, processing samples in the dark or triggering photorepair; Table 1). Within the studies, *Escherichia coli* has been used as the most common challenge organism (in 10/10 studies) but some studies employed other strains that are important for human health like *Staphylococcus aureus* (*28*). As UV disinfection has been researched for a long time and there are countless studies in the literature, however, bacterial species other than *E. coli* or relevant to human health are not commonly investigated. Experiments were run with both, exponential and stationary phase cultures (*21, 24*). Sometimes just growth conditions, but no direct information about growth phase is given (*27, 28*). Sensitivity of microorganisms to UV light might not only change with type of growth phase but also with the specific growth rate (*37*). Moreover, it should be noted that some studies used reactors in single pass mode whereas others used recirculation. By direct comparison of those two settings, differences were identified (*3*). Finally, whereas some studies gave details on sample processing, the information provided is often incomplete. Overall, this led to the conclusion that there is no standardized protocol for biodosimetry performed in lab-scale flow-through systems. Therefore, we developed a robust, reproducible procedure for the relevant biofilm forming bacterium *A. citratiphilum* taking into account key operational parameters that were previously not consistently used or recorded based on our literature research (i.e., sterilization of the skid, importance of the hydraulic characteristics of the used system, or the order of flow steps, photo repair and sample processing).

**Table 1.**
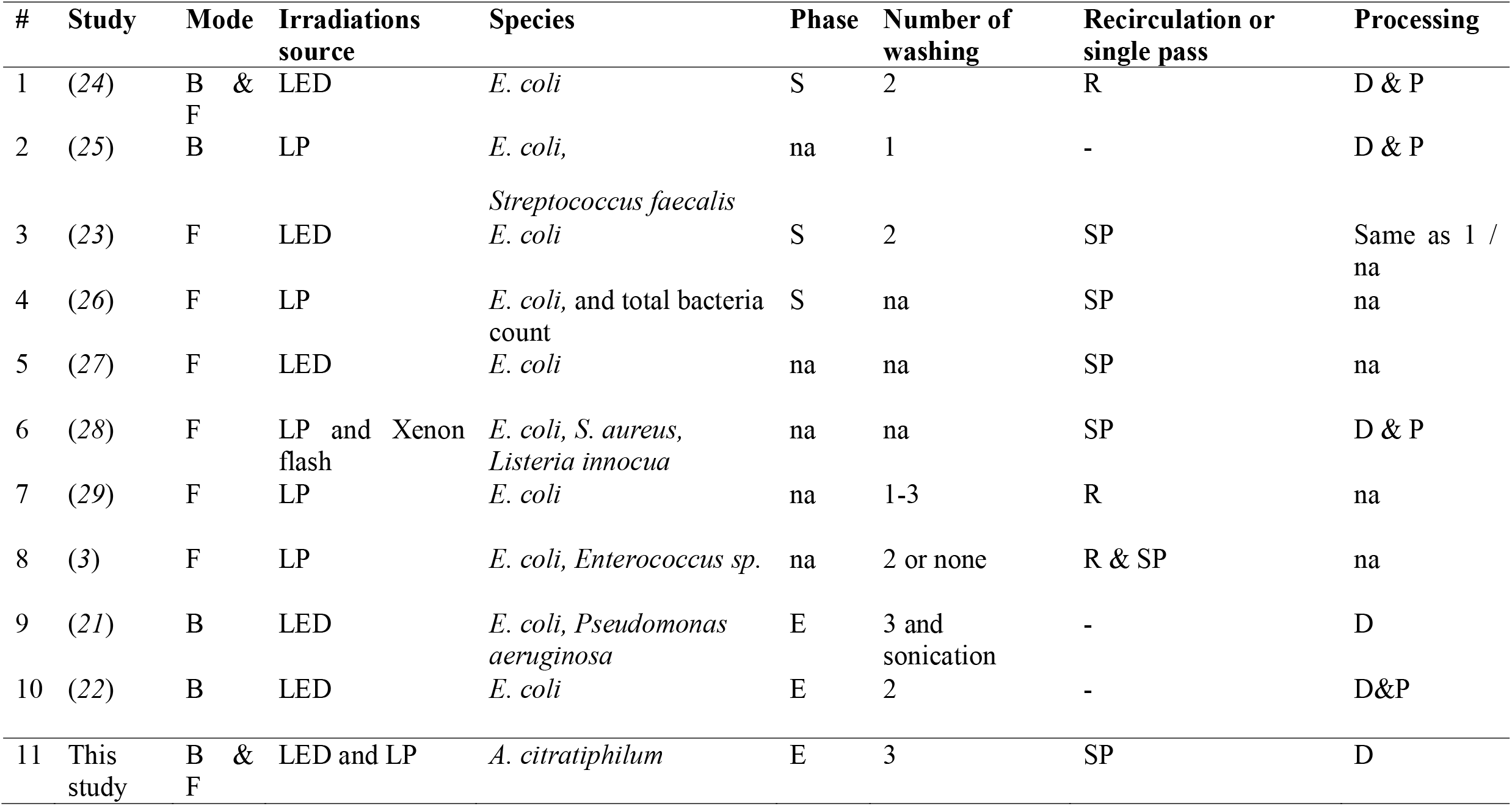
Overview on the methodology of selected studies, B = batch, F= flow through, S = stationary, E = exponential, R= recirculation, SP= single pass, D = dark, P = photorepair, na = information not available

### 3.1. Sterilization of experimental skid and contamination controls

Ensuring proper sterilization at the start of an experiment is critical, and appropriate controls are necessary to ensure a contamination free operation of the biodosimetry. Here, a method using 1% H_2_O_2_ was chosen, as the solution is easy to handle and compatible with most materials. To test if the sterilization by solely using H_2_O_2_ is sufficient, a negative control sample was taken during each experiment. The sample was taken during the flushing with sterile PBS and the LED switched off prior to the start of the experiment. Based on these observations it could be demonstrated that disinfection with 1 % H_2_O_2_ for 10 min is sufficient, as no CFU were found in the negative control samples.

### 3.2. Tracer test

The appropriate flushing feed volume required to achieve steady-state after changing a setting in a biodosimetry experiment was determined through two tracer tests (Figure 3). One tracer test was performed with a constant flow rate of 1.5 L h^-1^, one with higher flow rate of 6 L h^-1^. t_m_ for 1.5 L h^-1^ was 239.2 s and 76.5 s for 6.0 L h^-1^, respectively. The calculated *τ* for 1.5 L h^-1^ and 6.0 L h^-1^ were 289.5 and 72.1 s, respectively with a measured reactor volume of 120.3 mL. Whereas for 6.0 L h^-1^ space-time *τ* and mean residence time t_m_ were rather comparable, t_m_ for 1.5 L h^-1^ was smaller than *τ*, indicating presence of some dead space (*33*). Based on these results, it was decided to flush approximately 0.2 L, representing *Θ* ≈1.7, in between the different steps to achieve steady state. In addition, it should be noted, that the UV reactor is positioned in the last ∼40% of the skid. Hence, the 0.2 L were estimated to be four times the void volume from the UV reactor till effluent.

### 3.3. Order of flow steps

Based on the tracer test, three biodosimetry experiments with a driving current of 96 mA were performed for flow steps with 2.13, 2.83, 4.25, and 8.50 L h^-1^ resulting in approximately HRTs of 4, 3, 2, and 1 s, respectively (Figure 2). The sample order was random, but before the first sample, after sample number two, and after sample four, negative controls without irradiation were taken. Overall, the inactivation seemed not to follow the expected trend of increasing LRV with longer HRT. On average there could be still 300 to 5,900 CFU per mL detected in the samples of all HRTs, leading to the assumption that higher inactivation should still be possible. The reason for not seeing the expected trend of higher inactivation with higher HRT becomes more obvious when ordering the samples chronologically (ordering them as they were taken and by HRT) as shown in the supplementary material (Figure S3). After taking the blank samples (before samples one and three) the LRV is reduced.

**Figure 2.**
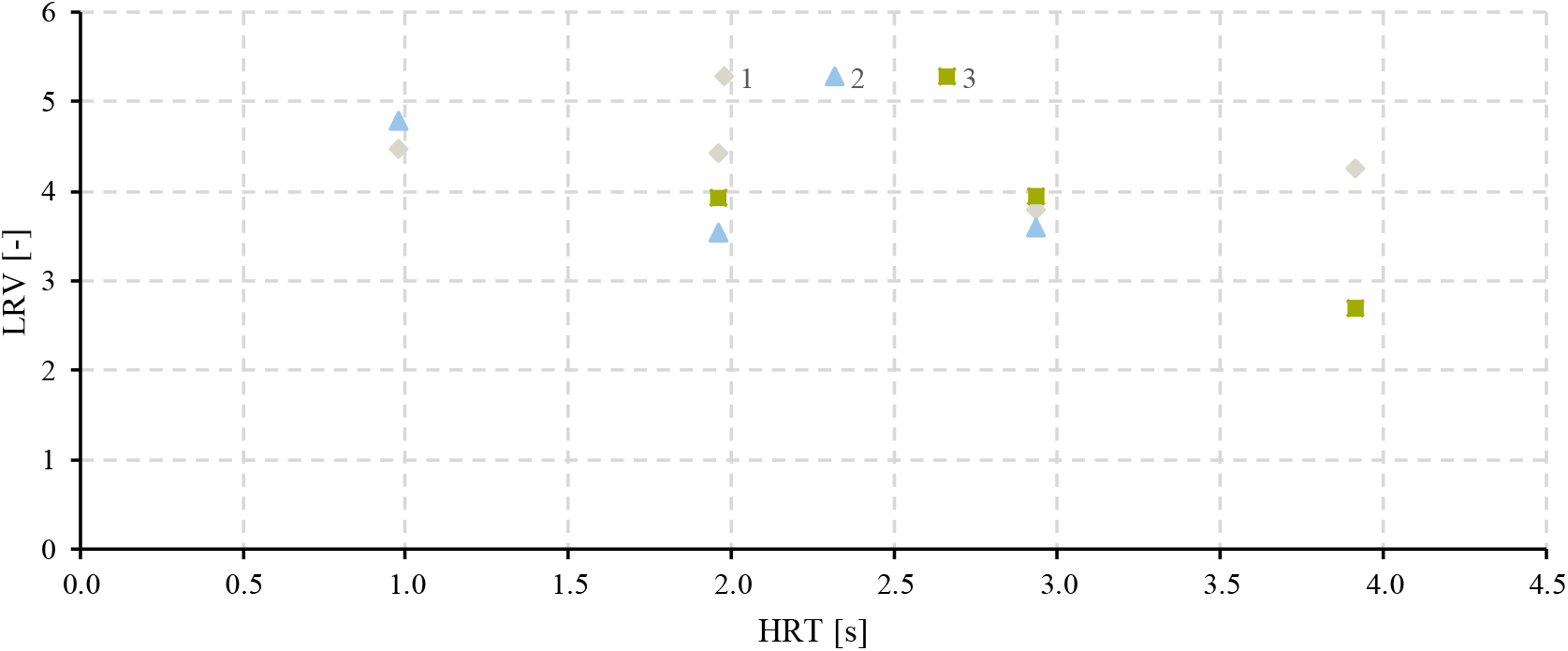
Logarithmic removal values (LRVs) based on biodosimetry using A. *citratiphilum* with a driving current of 96 mA over hydraulic retention time (HRT)

The reason for these inconsistencies in LRVs can be found when taking a closer look at the residence time distributions. Two models were fitted to the tracer curves to compare results to ideal reactors. The cumulative residence time distribution curve for the 6.0 L h^-1^ seems to fit with the CSTR. On the contrary, when looking at the curves of 1.5 L h^-1^, considerable differences can be seen (Figure 3). First, in comparison to the CSTR, the curve is shifted to a smaller *θ*. As discussed this was also seen as t_m_ < τ, leading to the assumption of dead spaces present (*33*). To compare the shape of the curves without offset, the F curve of 1.5 L h^-1^ of the tracer experiment was added with *E*(*θ*) = *t*_*m*_ · *E*(*t*) and 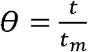 (Figure 3). Here a long tail out was present for *θ* > 1.5, however, as the used tracer was limited in precision with a maximal detectable change of 1 μS cm^-1^ for the conductivity sensor, the long tailing cannot be analyzed in more detail. The microbiological solution itself may serve as a step tracer to further resolve the long tail. Thereby system could be first filled with microbial solution and then flushed with sterile DI water to check which volume is needed to flush to have no CFU measurable anymore.

**Figure 3.**
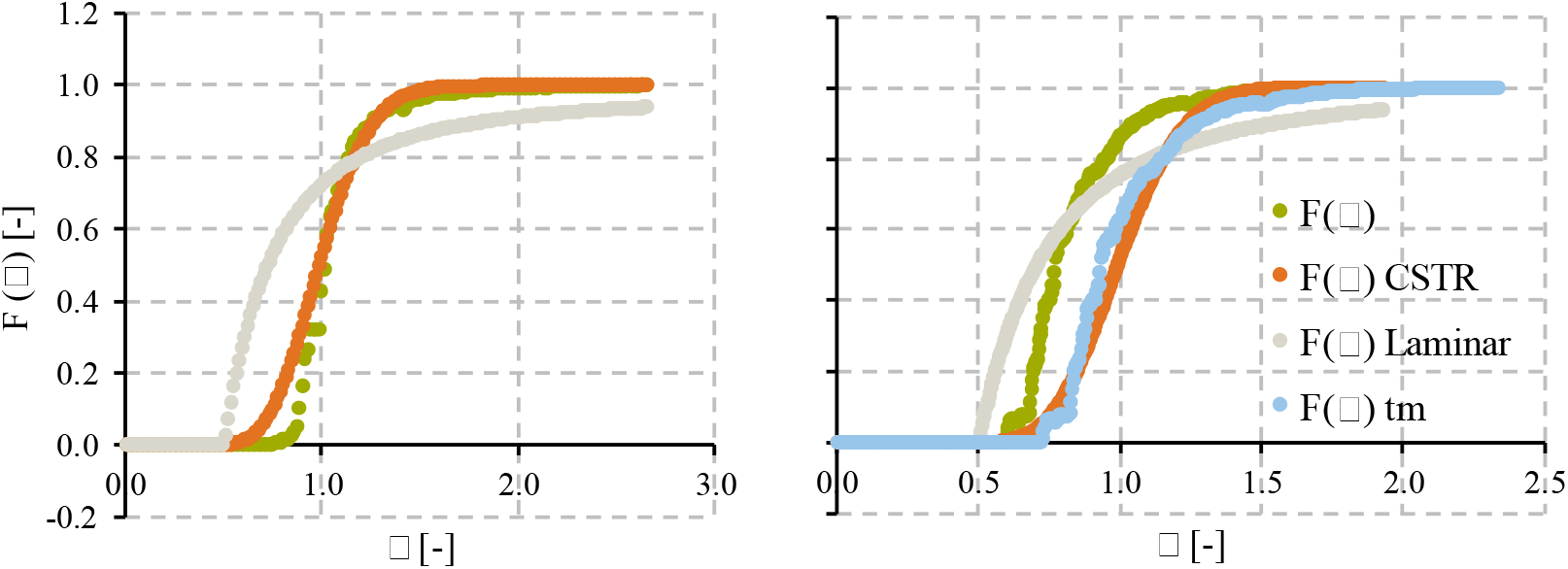
Fitted cumulative residence time distribution curves for different flow rates 6.0 L h^-1^ (left) and 1.5 L h^-1^ (right).

For the biodosimetry, the long tail can influence the inactivation of microorganisms: to confirm a LRV of 4 after a negative control reliable (with an error < one percent), the system needs to be flushed until an F(t) of 0.999999 is reached. Depending on the shape of the tail, this could increase the volume of microorganism solution to be flushed to unpractical volumes. As an extreme example the laminar flow model is considered with F(*θ*) = (1-1/(4 *θ*^2^)) for *θ* > 0.5 (*34*). *θ* required to reach 0.999 is 15.8 whereas for 0.9999 it is 50. So basically for pure laminar flow 50 times void volume flushing with sterile solution would be needed to reduce microbial count in the effluent from 10^6^ to 100 CFU mL^-1^.

One solution to reduce the void volume caused by the long tail of the residence time distribution is to reorder the flow steps, starting with the highest UV irradiation and ending with the negative controls. Assuming that due to flushing 0.1% of cells are still present in the samples of the next step, when running samples according to increasing cell numbers, the error is negligible. On the contrary, protracting 0.1% of cells from blank sample can have tremendous effects has it limits measurable LRV to ∼3 for the next sample. To confirm these assumptions, an experiment with a reduced driving current/fluence rate (24 instead of 96 mA) and with the new order of flow steps was performed. Thereby only an HRT of 1 s yielded in countable colonies in the sample. This leads to the assumption that the counts as shown in Figure 2 were only caused by the long tail of residence time. Following the new order was maintained and the driving current was further reduced to 16 mA. In addition, flow steps were changed to 1.06, 1.42, 2.13 and 4.25 L h^-1^ resulting in approximately 8, 6, 4, and 2 s HRT. The results of one representative experiment are shown in Figure 4. Here the expected trend of increased inactivation with rising HRT became noticeable even though in the step of 8 s less than 25 CFU were present and therefore results were not considered.

**Figure 4.**
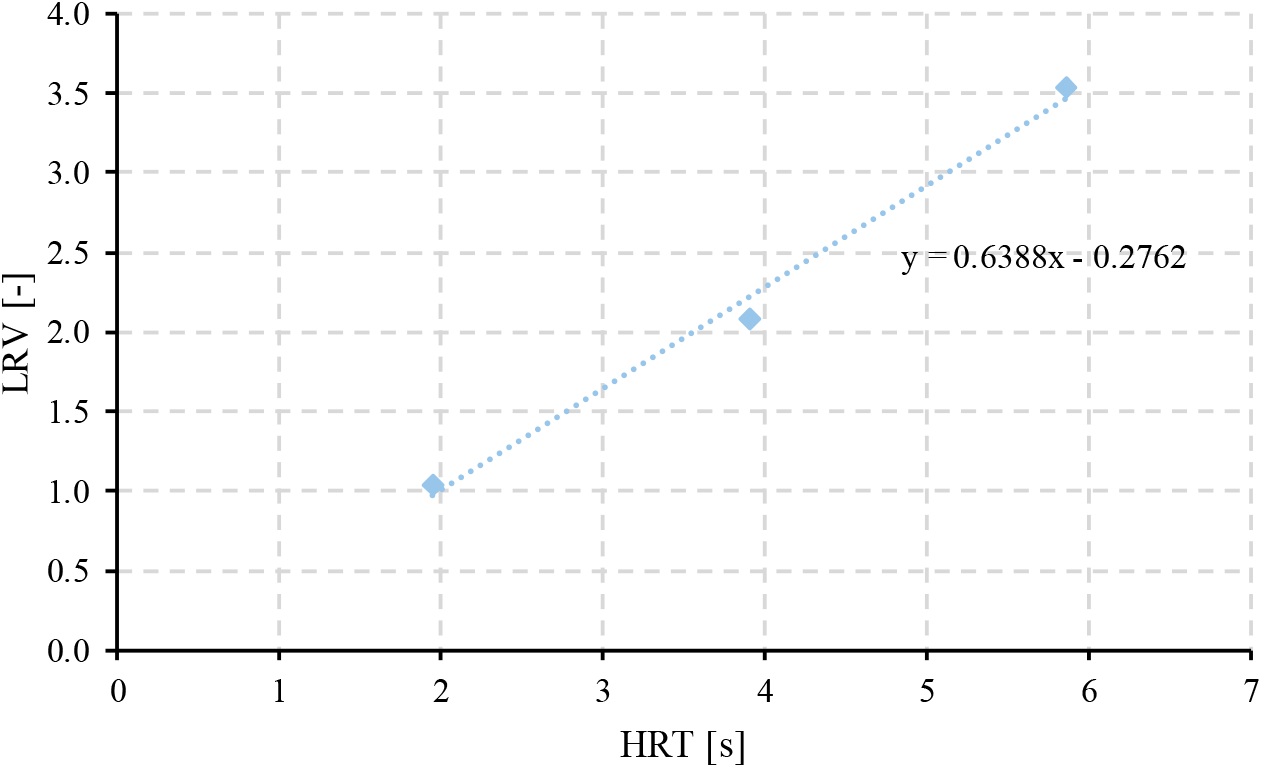
Biodosimetry experiment with a driving current of 16 mA and samples ordered with decreasing fluence.

On top of the discussed effects of long tails, adsorption-desorption effects could further influence the results. To avoid errors, it is therefore necessary to take the samples from the highest LRV/fluence in decreasing order. This strategy stands in accordance with NSF/ANSI 55 (*18*) according to which the UV experiment needs to be executed before the sample without UV disinfection. Alternatively, for experimental skids described in the U.S. EPA guidelines (*17*) influent samples are taken from a separate sampling point before the UV reactor, which can avoid protraction. However, taking an influent sample before the reactor might not be practical in the combination of treatments and does not reduce errors between different UV steps.

### 3.4. Photo repair and adhesion to plastic labware

Despite the improvement of results with the enhanced order of flow steps, it was recognized that samples sometimes showed inconsistencies between serial dilutions. This effect was especially present for undiluted samples and dilution 10^−1^. As an example, for an undiluted plated sample 253 CFU could be counted, whereas there were only 5 CFU present in the 10^−1^ dilution. For this inconsistency, two possible reasons were suspected and tested: photo repair, and adsorption to plastic labware.

#### 3.4.1. Photo repair

Under exposure to near UV light (310 – 480 nm) photo repair of DNA damages can occur (*1, 7*). When Oguma et al. (*7*) irradiated 254 nm UV treated samples of *E. coli* with fluorescent tubes with 18 W (0.1 mW cm^-2^ at sample surface for 360 nm) for 3 h, they observed recovery from approximately 3 LRV to 0.9 LRV. The sterile working bench used in this study was equipped with two 36 W fluorescent tubes. As processing of all samples usually took considerably longer than 1 h, photo repair during sample preparation might be significant, as photo repair was suspected to be rather limited by time than by irradiance (*1, 7*).

The photo repair effect might be further increased as samples were taken in transparent 15 mL tubes, while 1.5 mL tubes were used for serial dilutions. The 15 mL tubes are suspected to provide a larger surface and probably also higher transmission for visible light (Figure S4). This could lead to discrepancies between the CFU in undiluted and diluted samples. To reduce the effects of photo repair, the undiluted samples were taken in 1.5 mL tubes and stored on ice in the dark until the cell spreading process was conducted. The laboratory itself was darkened and commercially available red filters were installed in the sterile working bench (the only source of light in the lab). Even though the effects of photo repair are well known, methodological details are often missing (Table 1). To avoid artifacts of photo repair, this is however important information that has to be reported accordingly.

When comparing results with the red filter installed (Figure 5) to the previous outcome (Figure 4), LRVs are increased approximately by 1, which is likely attributed to avoided photo repair. Furthermore, without using the red-light filters, a shoulder (negative intercept), typical for repair processes was observed, which is not the case anymore.

**Figure 5.**
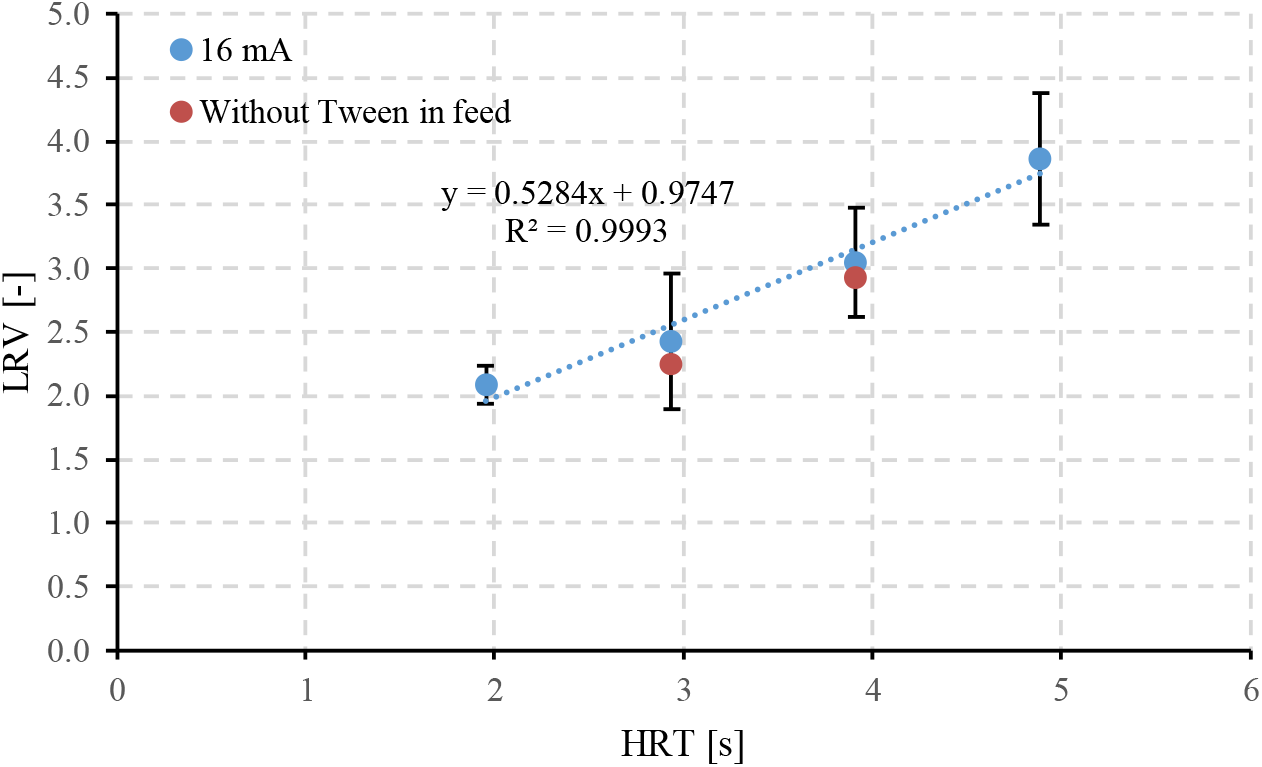
Final biodosimetry results at 16 mA driving current and without Tween20 added to the feed

#### 3.4.2. Adsorption to plastic labware

In addition to the optimized sample processing method and the installation of the red filters, the adsorption on plastic labware was examined, as it was previously shown that this may influence results (*38*). Therefore, instead of standard Eppendorf tubes, different low-binding protein or low-binding DNA tubes were used together with low retention tips for pipetting. Similarly as in the study of Richter et al. (*38*), differences between different brands of low binding tubes were found. As an example, for one tube, there were 22 CFU observed at a dilution of 10^−1^ whereas 206 CFU were counted for another tube. Although some artifacts remained with all tested tubes, testing different suppliers can improve inconsistencies in the CFU dilution series. Overall, Richter et al. (*38*) observed a correlation of phage adsorption to plastic labware based on hydrophobicity. More specific, they observed a threshold of 95° wetting angle below which tubes were marked as safe for adsorption. Tween 20, a common surfactant reducing protein adsorption, was found to reduce the wetting angle when used as a coating (*38–40*) and seems therefore promising in reducing bacterial adsorption as well. Following the recommendations of (*38*), 0.02 mL Tween20 were added per L PBS in the feed solution, the solution for dissolving washed *A. citratiphilum* pellets, and in the PBS used for dilutions.

Additionally, we considered a 95% confidence interval threshold that will mark and exclude samples with adsorption artifacts automatically. Basic cell counts N for each sample are assumed to follow a Poisson distribution, but it is not clear if cell counts after an inactivation experiment still follow Poisson distribution (*41*). Nevertheless, confidence intervals for Poisson distribution were calculated based on the Chi-square distribution 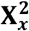 as following (*42–44*):

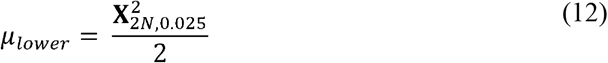

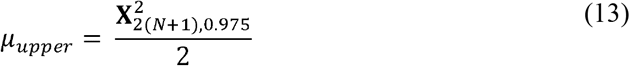

Hence if 10 · μ_upper_ of dilution 10^−x^ > μ_lower_ of dilution 10^−(x-1)^, the condition was fulfilled. This provides a possible, objective and easy implementable quality control step for biodosimetry experiments.

Following the improved procedure inconsistencies between dilution steps could not always be avoided, but were reduced. Results of the improved method can be seen in Figure 5. Hereby flow rates were additionally adapted to 1.7, 2.13, 2.83, and 4.25 L h^-1^ leading to HRTs of approximately 5, 4, 3, and 2 s. It is noteworthy that for one experiment less than 25 colonies were counted at the 5 s HRT step potentially leading to a higher deviation of the replicates.

To test if Tween20 in the feed and UV irradiation has additional disinfection effects on the cells, two HRTs were tested without Tween20 added to the feed. As those results fall within the standard deviation of the test with Tween20 in the feed, it is therefore assumed, that Tween20 for the used setting has no significant effect on the inactivation of *A. citratiphilum*. Nevertheless, this could change depending on the irradiation source and tested microorganism. Tween20 was found to serve as food source for *A. citratiphilum* (*30*) and therefore no toxic effects of Tween20 in the used concentration alone can be assumed.

### 3.5. Final procedure

Considering the above-discussed points, we subsequently developed a reproducible, standardized protocol, which is presented in details in the supplementary material (section 6). Preparations include the simple tracer test, sterilization procedure, cultivation, setting-up the growth curve and washing of *A. citratiphilum*, as well as preparing feed solution for experiments. The experimental part describes taking the negative control samples, order of flow steps, sampling procedure and controls necessary. Finally, processing of samples is defined including measures against photo repair and checking for adsorption on labware.

## 4. Conclusion

While biodosimetry with phages of batch systems is a highly standardized procedure, bacterial based biodosimetry protocols are not yet standardized, in particular for flow-through systems. Differences of methodology can be seen in the preparation of bacterial solutions including the growth phase and washing. The experiment itself varies between single pass and recirculation experiments and the order of flow steps is commonly not addressed. The processing of samples varies from avoiding to triggering photo repair or no information given at all. Within this study it was shown that the reactor hydraulics and possible long tails can heavily impact the results. To avoid and minimize those errors, it is recommended to take the samples descending with fluence. While processing of the samples possible photo repair and adsorption of cells to labware was observed. Even though, photo repair was not intended in the first place, environmental light might have influenced results during processing of the samples. Hence it is recommended to work only under red light to limit opportunities for photo repair. Installing red light filter covers for fluorescent tubes might be an easy solution for sterile working benches. Concerning adsorption to plastic labware, an objective criterion was presented, adsorption was reduced by the usage of low-binding tubes and Tween20, as well as Tween20 check for adverse effects. Finally, a hands-on protocol for using the indicator organism *A. citratiphilum* was proposed including a simple tracer test, sterilization of the skid, and the other critical points related to the order of sampling, photo repair and reducing cell adsorption. The proposed method is reproducible and effectively avoids artifacts. Considering the critical points during biodosimetry for microbial cells greatly increases reproducibility and transferability of results between studies.

## Corresponding Author

Christian Wurzbacher - Chair of Urban Water Systems Engineering, Technical University of Munich, Garching, 85748, Germany

## Author Contributions

The manuscript was written through contributions of all authors. All authors have given approval to the final version of the manuscript. Conceptualization, P.S. and C.W.; methodology, P.S., C.W. and M.K.; validation, P.S., C.W. and M.K..; formal analysis, P.S.; investigation, P.S. and M.K.; resources, J.E.D. and C.W.; data curation, P.S. and M.K; writing—original draft preparation, P.S.; writing—review and editing, J.E.D., C.W. and M.K.; visualization, P.S.; supervision, J.E.D. and C.W.; project administration, J.E.D. and P.S.; funding acquisition, C.W. and J.E.D.

## Notes

The authors declare no competing financial interest.

## ACKNOWLEDGMENT

The authors would like to express their sincere gratitude to the staff of the Technical University of Munich for supporting this work. Many thanks to UV-EL GmbH for providing the LED-reactor.

## ABBREVIATIONS

CBA: collimated beam apparatus
CFU: colony forming units
EC: electric conductivity
HRT: hydraulic retention time
LRV: log removal value
LP: low pressure
MP: medium pressure
OD: optical density
PBS: phosphate-buffered saline
RED: reduction equivalent dose
RTD: residence time distribution
UV: ultraviolet
UVT: ultraviolet transmissions.

