## Supplemental Material for "A practical bacterial biodosimetry procedure to assess performance of lab-scale flow-through ultraviolet water disinfection reactors"

### Supporting information: A practical biodosimetry procedure for lab-scale flow-through ultraviolet water disinfection reactors

*Philipp Sperle*[a], *Mohammad S. Khan*[a], *Jörg E. Drewes*[a*], *Christian Wurzbacher*[a]

[a] Chair of Urban Water Systems Engineering, Technical University of Munich, Garching, 85748, Germany

*

1. Schematic drawing of the utilized UV-LED reactor

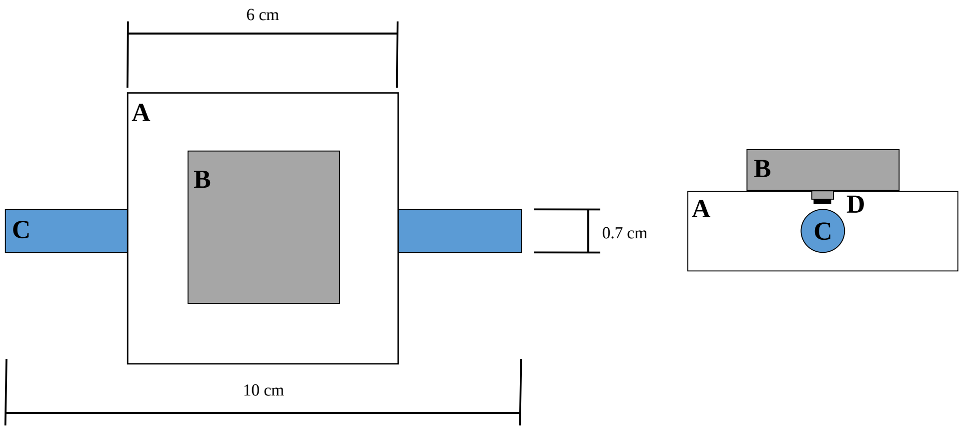


**Figure S1.** Simple schematic of the used UV-LED reactor; **A** reflective PTFE block, **B** LED cooling, **C** silica glass pipe, **D** LEDs; left: top view, right: sectional view

2. Optimized cultivation

To monitor cell growth, OD was checked over time for different cultivation approaches to find a practical procedure. With an iterative approach, nutrient concentrations and inoculation method was optimized. A single colony of a one-week-old streaked *Aquabacterium citratiphilum* plate was inoculated in 80 mL of bouillon in a 250 ml Erlenmeyer flask. By using twice the concentrated bouillon after an incubation of 48 h at 20 °C and shaking of 120 rpm, high cell concentrations in exponential growth phase were obtained.

3. Growth curve

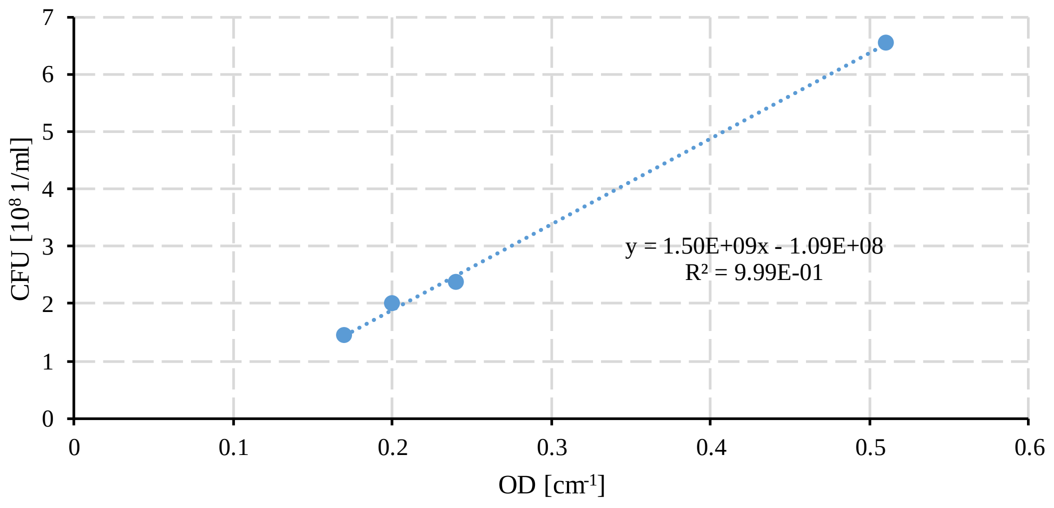


*Figure S2 Developed growth curve without low binding tubes and Tween20*

4. Results biodosimetry 96 mA, chronologically ordered with sampling

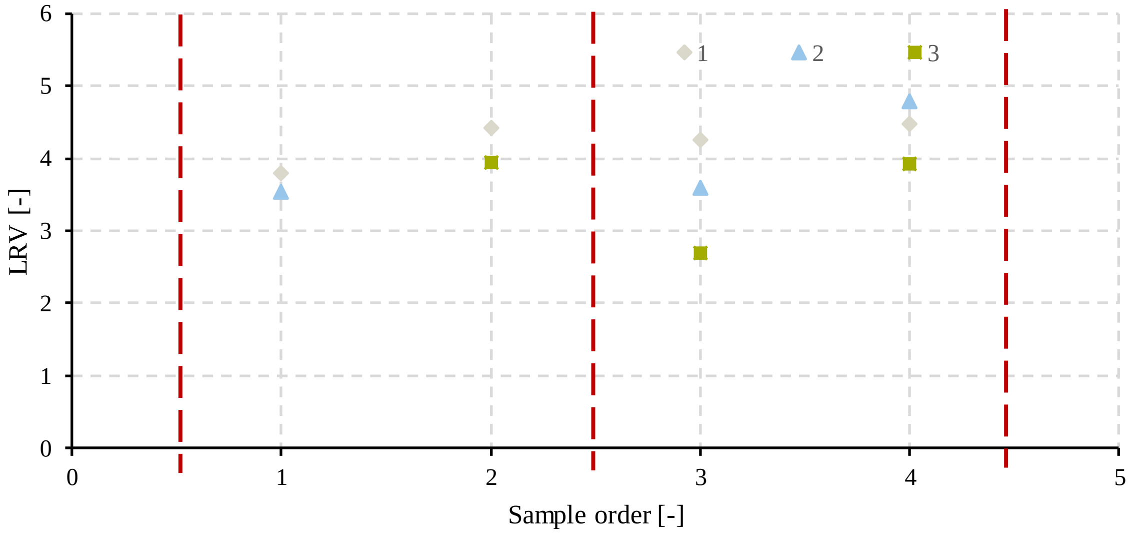


*Figure S3. Results biodosimetry 96 mA, chronologically ordered according to sampling; red lines representing the time when a blank sample was taken*

#### 5. Sampling tubes 15 and 1.5 mL

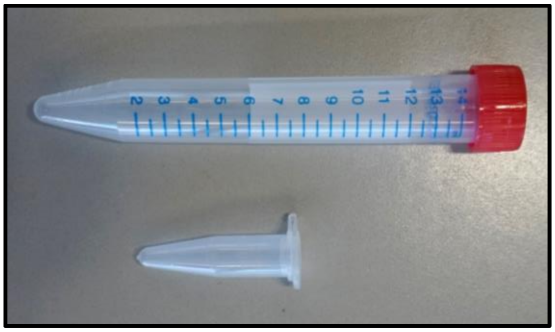

*Figure S4: Comparison of 15 and 1.5 mL sample tubes*

#### 6. Hands on protocol for simple tracer test

The protocol shown here was developed according to needs of the study and is based on standard methods for microbiology not further explained here. Depending on the needs, parts of the procedure might be adopted, bearing in mind the critical parts discussed. Please note that appropriate disposal of used chemicals is needed according to the local circumstances but not further described here.

#### Materials and Equipment

- Active *A. citratiphilum* (DSM 11900) streaked on an 830a agar plate
- Autoclave
- Bunsen-burner
- Cell density meter or apparatus to measure absorbance at 600 nm with disposable cuvette
- Centrifuge
- Colony counter
- Erlenmeyer flasks (250 mL)
- Incubator
- Inoculation loop
- Micropipettes with disposable, sterile (low retention) tips, at least 200 and 1.000 µL
- pH probe
- Pipette controller and disposable, sterile pipettes, e.g. 25 and 50 mL
- Portable electrical conductivity (EC) meter
- Spectrometer for measuring absorbance in irradiated wavelength with silica glass cuvette
- Sterile (low binding) 1.5 mL Eppendorf tubes
- Sterile 830a agar plates prepared according to (*1, 2*)
- Sterile 50 mL falcon tubes
- Three identical Drigalski spatulas
- Volumetric flasks, e.g. 250, 500, 1000 and 2000 mL

- Vortex mixer

#### Solutions

**Solution A – Deionized (DI) water**

Should show an (EC) < 10 µS $cm^{-1}$.

**Solution B – Tab water or alternative tracer solution**

Used for positive step tracer test, should show an (EC) > 500 µS $cm^{-1}$.

**Solution C – Sterile modified 830a boullion**

Prepared according to medium plates prepared according to (1, 2), but in double concentration and without agar.

**Solution D – Sterile phosphate buffered saline (PBS) plus Tween20**

1x 2.5 L, 2x 0.5L prepared according to NSF/ANSI 55 (*3*) added with 0.02 mL Tween20 per L, autoclaved and stored in a closed bottle. Add magnetic stirring bar for the 2.5L before autoclaving.

**Solution E – Ethanol**

80% ethanol for disinfection.

**Solution F – 1% $H_2O_2$ solution**

2x 0.25 L 1% H2O2 prepared form concentrated, for room temperature stabilized H2O2 stock solution on the day of experiment.

**Solution G – Sterile MilliQ water**

1L autoclaved MilliQ stored in a closed bottle.

**Solution H – Sterile phosphate buffered saline (PBS)**

2x 1 L prepared according to NSF/ANSI 55 (*3*), autoclaved and stored in a closed bottle.

**Preparations and experiment**

Determine the hydraulics of the system, steady state conditions, and the volume for flushing between samples. It should be performed at two flow rates covering the range of flow rates to be used in the biodosimetry experiments. The tracer test described below represents a simple solution that can be modified if needed with different tracer solutions.

1. Tracer test:
    - Determine space-time $\tau$ of the system by measuring and calculating void volume or alternatively first fill the system with DI water (solution A) and completely drain it again. Measure volume or weight of drained water to calculate space-time, e.g. 100 g of drained water at 20 °C represent ~100 mL, with a flow rate of 1 L $h^{-1}$ $\tau \approx$ 360 s.
    - Analyze EC tracer solutions (solutions A and B).
    - Connect conductivity meter to the outflow.
    - Fill system with DI water (solution A).
    - Stop pump and switch to tracer solution (tap water (solution B)).
    - Start pump at desired flow speed, e.g. 6 L $h^{-1}$ and continuously monitor EC till it becomes steady.
    - Calculate mean residence time as described in Chapter 2.
    - Check cumulative residence time distribution for volume to be flushed to reach steady state, F(t) at least >0.97, ideally >0.99.

Next to the knowledge on the hydraulics, key parameters (growth conditions and growth curves) for the challenge organism has to be setup, i.e. here for *A. citratiphilum.*

2. Growth curve:

  - Inoculate an active distinct colony of *A. citratiphilum* (a colony grown at 20 °C on 830a agar plates after streaking seven days ago using three-phase streaking pattern or similarly) in desired amount (e.g. 80 mL) of bouillon (solution C) in a 250 mL Erlenmeyer flask and incubate at 20°C and 120 rpm. If time for cell growth needs to be increased, several colonies may be inoculated.
  - Take samples in regular (e.g. 8 h) or practical intervals and measure optical density (OD) using the Cell density meter.
  - When taking OD reading first measure and the subtract blank reading of boullion (solution C).
  - Equilibrate 830a agar plates to room temperature if stored at 4 °C.
  - Prepare 10-fold dilutions using PBS with Tween20 (solution D) in 1.5 mL Eppendorf vials and plate 0.1 mL of the vortexed 10-fold dilutions on the 830a agar plates.
  - Use the Drigalski spatulas for streaking till a resistance between spatula and plate can be felt. Disinfect spatula first with 80% ethanol (solution E) and then flame it using a Bunsen-burner. Let Drigalski spatula cool down and use other sterile Drigalski spatula meanwhile.
  - Count colonies after plates been inverted and stored in the dark for seven days at 20 °C.
  - Count colonies using a colony counter.

- Build linear regression curve of OD vs. CFU $mL^{-1}$, considering only plates with 25-250 colonies analogous to (*3*).
- Continue to check OD till it reaches a maximum to investigate when stagnate phase is reached (e.g. > 0.6 $cm^{-1}$).

To avoid contaminations and false positive signals, several precautious disinfections steps are required before and after every experiment.

3. Disinfection procedure:
    - Freshly prepare two times the 1% $H_2O_2$ solution F in a volume bigger than the systems void volume (e.g. 250 mL, when void volume around 120 mL).
    - Flush system with DI water (solution A).
    - Switch feed to $H_2O_2$ (solution F) and discard the first void volume.
    - Start recirculation for 10 min at flow speed similar as used in the experiments or slightly higher, e.g. 10 L $h^{-1}$.
    - Sterilely connect MilliQ water (solution G) as feed and discard void volume.
    - Disinfect the outsides if influent and effluent pipes if necessary using surfactant, e.g. 80% ethanol (solution E) or heat.
    - Collect $H_2O_2$ and discard according to your local regulations.

Depending on the time *A. citratiphilum* need to be cultivated to reach the desired cell number, in our case around 1.25 $10^{10}$ CFU (typically reached after ~48 h), starting time to prepare the feed solution needs to be adopted.

4. Preparing *A. citratiphilum* feed solution:
    - As described in section 2 (growth curve) inoculate an active colony in 80 mL bouillon in an 250 Erlenmeyer flask and inoculate at 20°C with 120 rpm. I can be

practical to prepare two of this cell cultures, to avoid waiting periods on the day of experiment.

- After approx. 48 h, check OD.
- Calculate cell number according to growth curve.
- Check if sufficient cell number (~1.25 $10^{10}$ CFU) is reached (in our case within the OD the range of 0.3 to 0.5 $cm^{-1}$).
- When cell number is sufficient and cells are still in exponential growth phase e.g. OD <0.6, start washing procedure.
- For washing fill cell suspension in sterile 50 mL falcon tubes and centrifuge at 5000 rpm at 4°C for 10 min.
- Remove supernatant, resuspend cell pellet in sterile PBS (solution H) and vortex for 10 s.
- Repeat washing so in total suspension was centrifuged three times.
- For the last resuspension step use PBS plus Tween20 added (solution D) instead of only PBS.
- Depending on cell density, add needed volume of washed cell suspension to the 2.5L of sterile PBS plus Tween20 (solution H) that shall be used as feed and shake it. Make sure before the PBS was sterilized a magnetic stirring bar was added if needed.
- It is important to start the experiment (5. Biodosimetry experiment) as soon as possible after the preparation to avoid cells adapting to new low nutrient condition (*4*), in our case <1 h.

To safe time, it is recommended to prepare low binding vials with 0.9 mL of PBS + Tween20 (solution D) that can be used for preparing 10-fold dilutions the day before the experiment. Further if agar plates are stored at 4°C, these should be brought to room temperature a couple of hours before the processing of the samples starts.

5. Biodosimetry experiment:

   - Connect 1L sterilized PBS (solution H) as feed to the system.
   - Take a negative control sample in the same vials used for sampling later (to prove that system is disinfected/ CFU free).
   - Turn UV source and let it heat up if needed (we used 10 minutes for our LED to reach steady conditions in the used temperature cabinet).
   - Set slowest flow rate through the reactor, e.g. 1.7 L $h^{-1}$.
   - Switch feed to *A. citratiphilum* solution prepared according to section 4.
   - Starting with slowest flow rate/highest fluence, e.g. 1.7 L $h^{-1}$ and 16 mA driving current for the LED.
   - As soon as flow stabilized, stop time to flush enough volume through the system to reach steady state (as calculated by tracer experiment, e.g. 0.2 L).
   - Take replicated samples in low binding tubes.
   - Immediately store samples cool and in dark (e.g. in a cooling box and on ice).
   - Switch to next flow step (descending with fluence) and repeat, e.g. 2.13 L $h^{-1}$ after 1.7 etc.
   - As last sample type, take control blank samples without UV (use different flow rates to ensure pumping speed is not affecting cell viability, e.g. 4.25 L $h^{-1}$ and 1.7 L $h^{-1}$).

After experiment is finished, samples should be processed as soon as possible. Please note that when e.g. the bacterial solution is transported over longer distances, a trip control as described by U.S. EPA (*4*) might be needed. Further, in case samples without UV are collection before the UV reactor, at least once a sample after the reactor without UV should be compared (reactor controls (*4*)).

6. Sample processing

   - All processing of the samples need to occur in a red light only laboratory to avoid photo repair effects.
   - Prepare 10-fold dilutions using low retention tips, low binding vials and PBS with Tween20 added (solution D).
   - Plate 0.1-0.15 mL on an agar plate, samples should to be processed in random order.
   - Use sterile Dirgalski spatula to spread sample on agar plate as stand procedures in microbiology till you feel a resistance (see section 2 for preparation of growth curve).
   - The following contamination controls are plated: Bouillion (solution C), PBS+Tween (solution D) used for dilution, PBS+Tween (solution D) used for solution (last washing step), PBS used for washing (solution H) and negative control taken before the start of experiment.
   - After plating, agar plates are inverted and stored in the dark for seven days at 20°C.
   - After seven days CFU can be counted using a cell counter (only consider 25-250 CFU per plate analogous to (*3*).
   - Check if inconsistencies among the sample dilutions are occurring (using equations 14 & 15).

- Calculate LRV by using geometrical mean of replicate samples (*3*).
